# The Phylotranscriptomic Hourglass Pattern in Fungi: An Updated Model

**DOI:** 10.1101/2022.07.14.500038

**Authors:** Yichun Xie, Hoi Shan Kwan, Po Lam Chan, Wen Jie Wu, Jiachi Chiou, Jinhui Chang

## Abstract

The “developmental hourglass” describes the morphological convergence to a conserved form at mid-stages of animal embryogenesis. The molecular hourglass pattern during embryogenesis was also identified across kingdoms. Previously, we reported young fruiting body as the conserved “waist” in mushroom-forming “developmental hourglass”. However, its robustness is doubted because of the fungal diversity. Additionally, fungi lack embryogenesis, and develop directly from spore to hyphae with morphological similarities during the transition. Here, we updated the “developmental hourglass” model in the life cycle of fungi, namely, spore germination, vegetative growth, and sexual reproduction. Germinating spores, both sexual and asexual, showed the strongest transcriptomic conservation signals across the phyla Mucoromycota, Ascomycota and Basidiomycota. Cross kingdom comparisons revealed high expression levels of “information storage and processing” genes at the waist stages of embryonic and non-embryonic developments in animals, plants, and fungi. The “developmental hourglass” might reflect the mutual transcriptome switches on developmental transitions in eukaryotes that are additional to embryonic organogenesis.

**Highlights:**

- Updated fungal molecular “developmental hourglass” model in the life cycle of fungi
- Germinating spores are the evolutionarily conserved “waist” across fungal phyla
- High expression levels of “information storage and processing” genes at the waist stages in the embryonic and non-embryonic hourglasses across kingdoms
- “Developmental hourglass” may reflect the mutual transcriptome switches on developmental transitions in eukaryotes

---

The morphological similarities in embryogenesis have long been discussed since von Baer’s laws of embryology were first proposed in nineteenth century (von Baer 1828). Embryos from various taxa appear to be very different at early embryogenesis; however, they converge to share more similarities at mid-stage of the embryonic development, and then diverge again at late embryogenesis and toward adulthood. Such pattern was known as the “developmental hourglass” (Duboule 1994; Raff 1996). The molecular “developmental” hourglass has been confirmed by transcriptome-based analyses, that demonstrate the high expression of evolutionarily older and conserved genes in the mid-embryonic stage across species in both plants and animals (Kalinka et al. 2010; Irie and Kuratani 2011; Levin et al. 2012; Quint et al. 2012; Drost et al. 2015; Hu et al. 2017).

For mushrooms, the early development of fruiting bodies looks like an embryonic process (Moore 1998). Our previous study has uncovered the hourglass pattern during mushroom formation in *Coprinopsis cinerea*. Young fruiting body is the developmental waist of fruiting, where meiosis, sporulation and stipe elongation took place sequentially (Cheng et al. 2015). Similar to flowering plants (Quint et al. 2012; Drost et al. 2015), the waist appears to be not exactly correlated to organogenesis as in animals. Transcriptome analyses on plant seed germination and floral transition have shown that post-embryonic hourglass patterns may come from regulations in developmental transition check-points, and decouple from body planning (Drost et al. 2016).

Unlike many other organisms, fungi lack a embryonic stage and develop directly from a single cell spore to hyphae in a clonal manner (Moore 1998). Filamentous fungi transform from spore to filamentous form by germination. The advanced, differentiated multicellular sexual fruiting bodies form after germination and hyphal aggregation (Kües 2000; Wang et al. 2014). Although fungal spores of various taxa are highly diversified in terms of their morphological characteristics and genetic background (sexual or asexual), the germination processes show morphological similarities and can be divided into several stages: dormant spore (the resting state), swelling (water uptake and size enlarged), polar growth (polarisome component recruitment and germ tube emergence), germ tube elongation (long axis doubling and hyphal tip elongation), and hyphal branching (the formation of multinucleate side branches) (Kües 2000; Osherov and May 2001; Sephton-Clark and Voelz 2018). Taking spore as the starting point of a new life cycle, whether the germination process of different fungi shares common evolutionary features like the embryogenesis in other kingdoms remains unclear.

Genomic and transcriptomic resources of fungi and other kingdoms have massively increased in the past few years. Many comparative studies have been conducted to clarify the driven factors in fungal morphogenesis and multicellularity construction (Kiss et al. 2019; Krizsán et al. 2019; Varga et al. 2019; Merényi et al. 2020; Miyauchi et al. 2020; Virágh et al. 2022). Questions have been raised on the robustness of the hourglass model in fruiting body development in some fungal clades (Merényi et al. 2022).

In this study, we tested the phylotranscriptomic patterns in fungi, using the indication of gene ages and transcriptome age index (TAI) profile, and sequence divergences and transcriptome divergence index (TDI) profile. The transcriptome profiles of the complete life cycle of *Coprinopsis cinerea* (Fig. S1) (Lau et al. 2020; Xie et al. 2020; Xie et al. 2021) and *Fusarium graminearum* (Kim et al. 2018), the representative species of the phyla Basidiomycota and Ascomycota, respectively, from sexual or asexual spore germination to vegetative growth and to sexual reproduction, were acquired and analysed. Additionally, the germination process of *Rhizopus delemar* (Sephton-Clark et al. 2018), a Mucoromycota, was included in the examination. We aimed to (i) provide a more detailed developmental hourglass model in fungi; (ii) reveal the similarities and differences on gene expressions and functional features during the developmental process in fungal species, and across the kingdoms Animalia, Plantae, and Fungi.

## Results and Discussion

### The Molecular Hourglass Pattern in Fungal Development

The Transcriptome Age Index (TAI) measures the mean evolutionary age of the transcriptome. It is the weighted mean of gene ages (phylostrata) for expressing genes. The phylostratigraphy approach was used to determine and assign the phylostrata (Text S1 and Fig. S2) (Cheng et al. 2015; Drost et al. 2015). *C. cinerea* and *F. graminearum* showed the minimum value of TAI during spore germination (Fig. 1A and B), indicating an older transcriptome age and a larger number of evolutionarily old genes functioned in this process. We further tested an early diverging lineage species, *R. delemar* (Phylum Mucoromycota). The germination profile of *R. delemar* displayed lower TAI at the polar growth and germ tube elongation stages than the fresh dormant spores and germinated hyphal growth stages (Fig. 1C). All TAI profiles displayed the high-low-high hourglass pattern and showed statistical significance using the reductive hourglass test (*P* < 0.001).

**Fig. 1.**
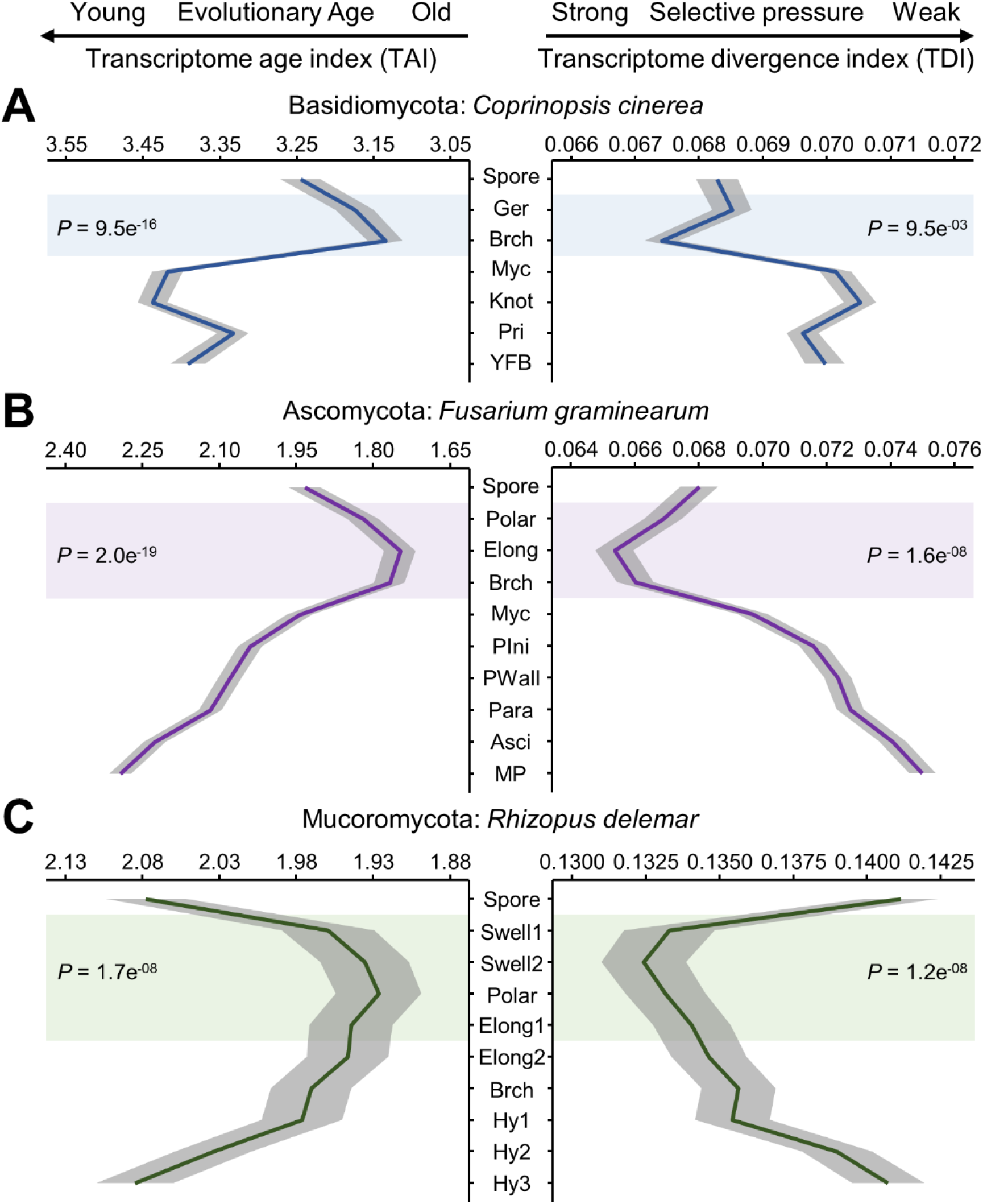
Phylotranscriptomic profiles during fungal development. (A) *C. cinerea, C. sclerotiger* as the TDI reference species; (B) *F. graminearum, F. fujikuroi* as the TDI reference species; (C) *R. delemar, R. arrhizus* as the TDI reference species. Gray area represents the standard deviation estimated by 1000 times of permutations. Colour shading indicates the predicted phylotypic period (waist stage) along the developmental process. P values are derived by application of the reductive hourglass test. Developmental stages are described in detail in Table S1-4.

Each phylostratum differentially contributed to the TAI profile. Old genes (PS1-2, orthologs shared across cellular organisms and eukaryotes) contained the majority of genes in all the three species and took up one-third to one-half of TAI (Fig. S3). They showed high relative expression during spore germination. The peak occurred earlier or later at the polar growth, germ tube elongation, or hyphal branching stages, which was a narrow interval of six to twelve hours as determined by the sampling time points. We suggested the existence of a highly conserved “waist” stage, but it was masked by the bulk-sequencing and weak synchrony of the spore germination culture. On the contrary, the expression peaks of young genes (PS3-10/12) were mainly found in the fruiting bodies and fresh spores, while the valley was observed at hyphal branching stages (Fig. S4 and S5). These results indicated the divergent gene usage and preferences of different developmental processes.

To further determine the regulation on either old genes or young genes drove the fluctuation of TAI, the mean relative expression levels of the phylostrata were computed (Quint et al. 2012). From fresh spores to hyphal branching stage, expression of young genes was down-regulated, while expression regulation on old genes was not significant; expression of young genes was later increased during sexual reproduction (Fig. S6). The old to young gene expression ratio maximised during germination in all three fungi. Furthermore, the number of expressed young genes decreased at the waist stage, while the transitions of old genes did not always consistent with the TAI profile (Fig. S7). These results suggested the young and divergent genes modulated the TAI profile by quantitatively decreasing the expression levels and number of expressed young genes (Quint et al. 2012).

Next, dN/dS ratio and TDI were computed to detect the recent evolutionary signals and selection pressure (Quint et al. 2012; Cheng et al. 2015). Reference species of different phylogenetic distance were obtained to minimise the bias (n≥3 at each taxonomy level). Most genes in all three species had a dN/dS ratio less than 1, indicating these genes were under purifying selection (Fig. 8-18 A). The TDI profiles displayed significant reductive hourglass patterns (*P* < 0.01), and the developmental “waist” occurred during spore germination as TAI (Fig. 1). The reductive hourglass patterns were robust across the reference species at different phylogenetic distances (*P* < 0.05, Fig. 8-18 B). The TDI profiles indicated that fungi were under different selective pressure during the development, and the hourglass was actively functioned and maintained (Cheng et al. 2015; Drost et al. 2015). Although sexual spore or asexual spores were used in different species, the hourglass patterns of TAI and TDI remained conserved for different spore types and genetic backgrounds. Thus, TAI and TDI profiles together revealed the high conservation of fungal spore germination across phyla and spore types.

Consistent with our previous study (Cheng et al. 2015), another waist-stage can be observed during the sexual reproduction of *C. cinerea*. Although the secondary waist was not observed in *F. graminearum* development, a smaller peak on old to young gene expression ratio occurred at the paraphyses stage, just as the peak showed at primordia stage of *C. cinerea* (Fig. S6B and C). In paraphyses or primordia, hymenia differentiate into basidia/asci and sterile cells, and the latter would support perithecia or fruiting body caps. Thus, cell differentiation and meiosis actively happen in those immature fruiting bodies (Kües 2000; Sikhakolli et al. 2012). In both *F. graminearum* and *C. cinerea*, the body plan was determined before these stages (Trail and Common 2000; Srivilai and Loutchanwo 2009), but the complicated multicellularity construction continued happening. Such observation was similar to the post-embryonic hourglass pattern during seed germination and floral transition in plants (Drost et al. 2016; Drost et al. 2017), and suggested the decoupling between hourglass pattern and body planning.

In *C. cinerea*, asexual reproduction and stress-resistant response are two alternative developmental paths. Oidia and sclerotia are produced in the above processes, respectively, and they function as the reproductive structures like the basidiospores formed during sexual reproduction. Their TAI and TDI were closed to vegetative growth and sexual reproduction, and higher than spore germination (Fig. S19 and S20), suggesting that these four processes evolved, adapted and fixed against various biotic and abiotic factors much later than the spore germination (Knapp et al. 2018; Kiss et al. 2019; Xie et al. 2020).

### Common Functional Regulations across Kingdoms

As the reductive hourglass pattern can be detected during spore germination in the three fungi from different phyla, we further sought to understand the functional features shared by these fungi, as well as the embryogenesis process of animals and plants (Drost et al. 2015). Developmental process was separated into early-, mid- and late-stage according to the high-low-high hourglass pattern, and genes were annotated and classified into Eukaryotic Orthologous Groups (KOG) (Fig. S21 and S22). Genes that followed the low-high-low expression pattern were enriched in informational KOG classes, and the high-low-high genes were enriched in metabolism (Fig. S23). On the genome-wide scale, at early-stage, RNA in dormant spores was inherited from the parent. Most activities were suppressed, except for a few KOG classes that were involved in metabolism and cellular processes and signalling.

Entering mid-stage, “information storage and processing” genes were actively functioned, and the expression peaks of these KOG classes were mainly reached during this period. At late-stage, the informational genes were down-regulated, while metabolic genes were highly expressed (Fig. S24-26). Similar regulatory patterns were also observed during the embryogenesis of *Drosophila melanogaster, Danio rerio* and *Arabidopsis thaliana* (Fig. S27-29). Genes of “translation, ribosomal structure and biogenesis” were highly expressed at the mid-stage in all six species (Fig. 2A). In brief, not only for fungi from different phyla, but also for animals and plants, high expression levels of informational genes and metabolic genes happened at the mid-stage (waist) and late-stage, respectively, while the expression of structural genes were fluctuated (Fig. 2B). The regulatory pattern on functional gene usage during the complex multicellularity formation could be conserved across three kingdoms, with or without the morphological similarity from single cell to multicellular structures (Quint et al. 2012; Cheng et al. 2015).

**Fig. 2.**
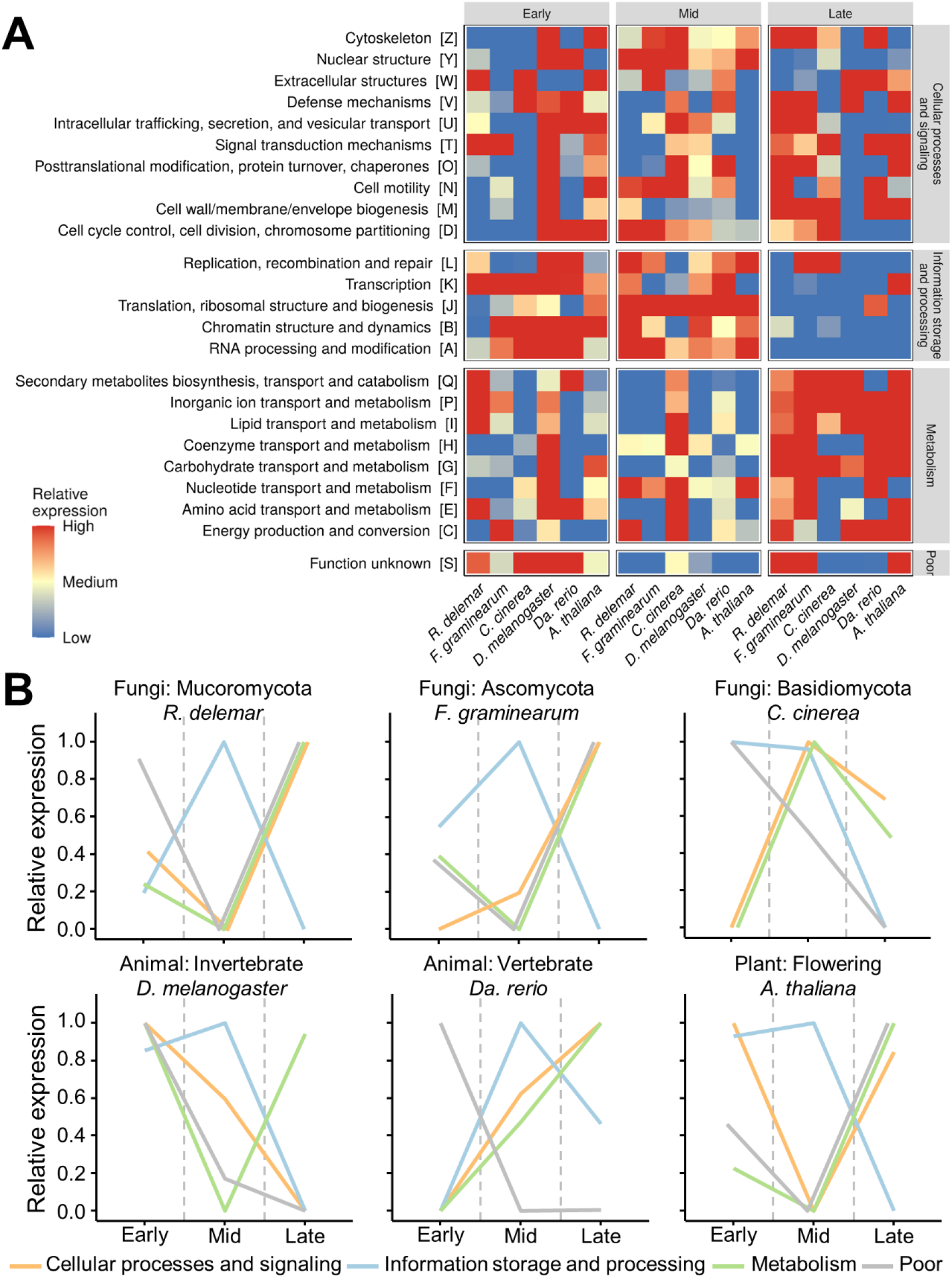
Common functional regulations across eukaryotic kingdoms. (A) Mean relative expression levels of genes of each KOG class; (B) Mean relative expression levels of each KOG ontology. Early-, mid-, and late-stages are defined according to the predicted phylotypic period. Relative expression levels of each stage are shown in Fig. S24-29.

We further asked if similar functional regulation happened in non-embryonic hourglasses. Therefore, we annotated the functional features of two post-embryonic hourglass processes, the seed germination and floral transition in *A. thaliana* (Drost et al. 2016). Again, informational genes were highly active at mid-stages, especially the “chromatin structures and dynamics” and “replication recombination and repair”, and were same to the functional classes found at the secondary “waist” stage in *C. cinerea* (Fig. S26, S30 and S31). These genes functioned in meiosis and mitosis of the above-mentioned developmental processes, and were associated with the increase in cell types and the number of cells on morphoanatomy. These features were conserved across kingdoms. We hypothesised that regulations of the fundamental processes during developmental transition could lead to the hourglass pattern, and accompany with the high expression of informational genes (Fig. 3). It was reported that the developmental chromatin accessibility might be associated with the transcriptomic hourglass of embryogenesis in vertebrates (Uesaka et al. 2022). But the effect of chromatin accessibility on the transcriptome divergence in other developmental processes remains unclear.

**Fig. 3.**
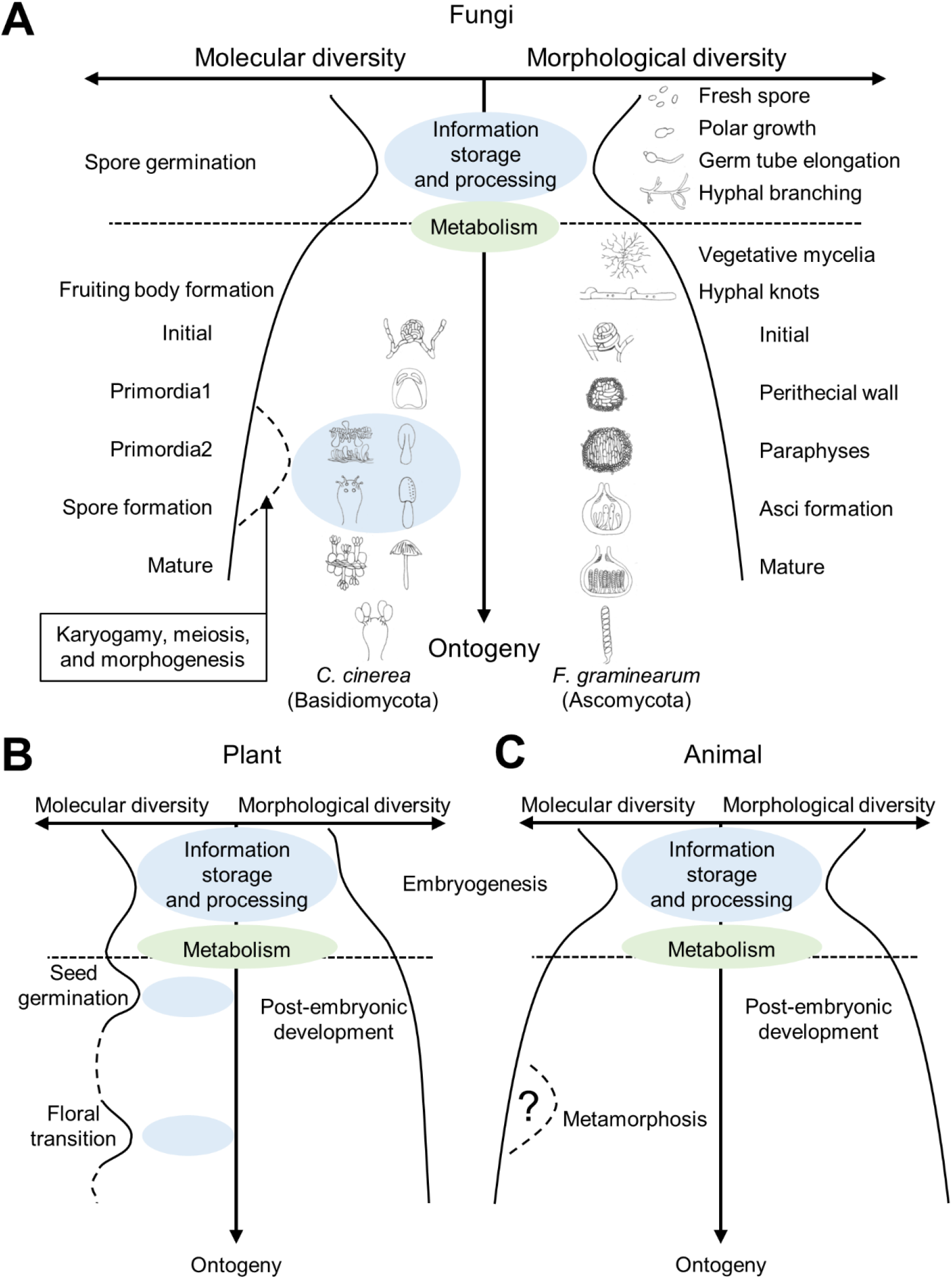
Common features on molecular and morphological developmental hourglass pattern across kingdoms. (A) Fungi; (B) Plant; (C) Animal. (B) and (C) are modified from Drost *et al*. (2017).

## Conclusions

The fungal kingdom has high morphological and genetical diversity. Our study modified the molecular “developmental hourglass” in fungi using the full life cycles and cross phyla comparisons. The spore germination process in different fungi shows morphological similarities and displays strong transcriptomic conservation signals. Functional regulatory similarities on both embryonic and non-embryonic developments suggested that the hourglass pattern could be a reflection of transcriptome switches in multicellularity construction across kingdoms, inclusive of but not limit to organogenesis and body planning. The developmental hourglass is observed in fungi.

## Author contributions

Conceptualisation: Y. Xie, J. Chang, H. S. Kwan; formal analysis: Y. Xie, J. Chang; software: Y. Xie; investigation: Y. Xie, J. Chang, P. L. Chan, W. J. Wu; writing: Y. Xie, J. Chang; funding acquisition: H. S. Kwan, J. Chang; supervision: J. Chang, H. S. Kwan, J. Chiou. All authors have read and agreed to the manuscript.

## Acknowledgement

We would like to thank Mr. Wenyan Nong, Dr. Xuanjin Cheng, and Dr. Amy Yuet Ting Lau for fruitful discussions, Ms. Jackie Wong for laboratory coordination and Ms. Josephine Leung for language editing.

## Funding

This work was supported by the General Research Fund (GRF 14116916 and GRF 14103817), Research Grants Council of HKSAR, PRC. This work was also supported by the Starting Fund (Starting Fund to J. Chang) from the Hong Kong Polytechnic University. The funders had no role in study design, data collection and interpretation, nor the decision to submit the work for publication.

## Data availability

Supplementary files, expression matrix, PS values, dNdS ratio, reference sequences, codes on bioinformatic analyses are distributed at https://github.com/xieyichun50/Fungal-hourglass.

## Notes

### Competing Interest Statement

The authors have declared no competing interest.

